# A method for faster purification of serine proteinases from *Bothrops alternatus* and *Bothrops moojeni* snake venoms

**DOI:** 10.1101/289991

**Authors:** M.A.G. Heleno, L.D. Santos, R.S. Ferreira, B Barraviera

## Abstract

Snake venoms are important sources of substances with a variety of pharmacological activities. Among the different proteins present in these venoms, snake venom serine proteinases (SVSPs) have important effects on the hemostatic system that influence the hemodynamic properties of blood. *Bothrops* genus snakes presented their venom richly composed of SVSPs thrombin-like, and the isolation of these enzymes is of great interest. In 1994, the Center for the Study of Venoms and Venomous Animals (CEVAP) - UNESP standardized the fibrin sealant derived from snake venom, replacing the bovine thrombin by gyroxin thrombin-like enzyme from *Crotalus durissus terrificus* (Rattlesnake) and human plasma fibrinogen by buffaloes cryoprecipitate. Despite chromatographic techniques for the purification of gyroxin be well grounded in the literature, that income is considered low. Thus, in addition to gyroxin, other thrombin-like enzymes could be employed in the composition of the new fibrin sealant after being standardized to the purifying and chromatographic performance and widely evaluated for biological activities. Therefore, it is extremely important that in our lab is deployed, standardized and validated a method for the chromatographic purification of other thrombin-like enzymes such as found in Bothrops snake venoms. Thus a two-step chromatographic procedure was developed to routinely purify serine proteinases from *Bothrops alternatus* and *B. moojeni* snakes venoms to provide new enzymes for improving the CEVAP’s heterologous fibrin sealant.

## INTRODUCTION

The *Bothrops* snake venoms contain a large variety of proteins and peptides affecting the hemostatic system. These proteins are classified as coagulant, anticoagulant or fibrinolytic factors (Iyaniwura (1991), Marsh (1994). The high selectivity of these snake venom proteins for especific blood coagulation factors makes these componentes potentially helpful tools to study the mechanisms of action, the regulation and the structure-function relationships of coagulation factors. Among these proteins, serine proteinases serve as tools to study molecular details in the activation of specific factors involved in coagulation and fibrinolytic cascades and are useful in treating various thrombotic and hemostatic conditions. Serine proteinases are found in microorganisms (Siezen, 1999), plants (Rudenskaya, 1995) and numerous animals (Neurath, 1984 e 1985). Snake venom proteases, in addition to their contribution to the digestion of the prey, affect various physiological functions. They affect platelet aggregation, blood coagulation, fi brinolysis, complement system, blood pressure and nervous system (Kini. 2005).

These enzymes share many biochemical and structural properties such as a conserved catalytic triad (Ser195, His57, Asp102), and its three-dimensional structure is highly conserved (Greer, 1990; Perona, 1995). Their structure, taken as a whole, is made of two β-barrels constituted of six-strands and separated by the catalytic residues, and of a C-terminal helical segment. Snake venom serine proteinases (SVSPs) found in *Bothrops* snake venoms are functionally similar to endogenous blood clotting enzymes. They interfere with maintenance and regulation of the blood coagulation cascade by cleaving specific bonds and activating proteins involved in blood coagulation, platelet aggregation, fibrinolysis and in the proteolytic degradation of cells resulting in an imbalance of the hemostatic system (Kini, 2005; Serrano and Maroun, 2005). Many other SVSPs convert fibrinogen into fibrin by cleaving fibrinopeptides A and/or B. As this activity resembles the activity of thrombin, these venom components are commonly named “thrombin-like” enzymes. Snake Venom Serine Proteinases are encountered in the venoms of several *Bothrops* species. The amino acid sequence homology shared between the SVSPs mentioned above is approximately 65%, however, the homology exhibited by these enzymes with mammalian serine proteinases such as thrombin and trypsin, ranges from 30% to 40%. (Costa *et* al., 2013)

The mechanism of action of MSP1 (*Bothrops moojeni*), cerastobin (*Cerastes vipera*), cerastocytin and cerastotin (*Cerastes cerastes*) on platelet activation is unknown (Marrakchi *et* al., 1995; Serrano *et* al.,1995). Bothropase and Reptilase are thrombin-like serine proteinase from *Bothrops* snake venoms and their administration *in vivo* results in a transient hypercoagulable state. They have been used to treat patients with urinary or gastrointestinal bleedings. Ancrod and batroxobin, two other serine proteinases from *Bothrops atrox* venom, have a fibrinogenolytic action and have been used in the treatment of deep vein thrombosis and peripheral vascular occlusive diseases.

In 1994, the Center for the Study of Venoms and Venomous Animals (CEVAP) - UNESP standardized the fibrin sealant derived from snake venom, replacing the bovine thrombin by thrombin-like enzyme from *Crotalus durissus terrificus* gyroxin, and human plasma fibrinogen by buffaloes cryoprecipitate. Despite chromatographic techniques for the purification of gyroxin be well grounded in the literature, that income is considered low. Thus, in addition to gyroxin, other thrombin-like enzymes could be employed in the composition of the new fibrin sealant after being standardized to the purifying and chromatographic performance and widely evaluated for biological activities. Therefore, it is extremely important that in our lab be deployed, standardized and validated a method for the chromatographic purification of other thrombin-like enzymes such as found in *Bothrops* snake venoms.

## PURPOSE

The purpose of this work was to certify a faster methodology than usual to isolate TLEs from *Bothrops alternatus* and *B. moojeni* snake venoms, carachterize them and compare with other SVSP TLE deposited in databases, aiming to obtain new enzymes for improving CEVAP’s heterologous fibrin sealant.

## METHODS

The venoms were obtained from the “pool” of venoms extracted from adult *Bothrops* snakes (*Bothrops alternatus and Bothrops moojeni*), of both sexes, individually microchipping, created and maintained in the CEVAP’s Serpentarium, located at Fazenda Experimental Lageado, UNESP, Botucatu. The venom extraction was performed according to the methodology developed by the technical team of the CEVAP ‘s Venoms Extraction Lab, micro-filtered, lyophilized, aliquoted, identified and stored in a freezer at −20 ° C until use. The venoms were fractionated by affinity chromatography on Benzamidine-Sepharose 6B and fractions representing peak 3 from the affinity chromatography were desalted and applied at a flow rate of 1 mL/min onto reversed-phase liquid chromatography (RP-HPLC) C18 column (20 × 250 mm). The eletrophoretic analysis for evaluation of purity and determination of relative molecular mass (Mr) was performed in SDS-PAGE under denaturing conditions using as molecular weight standards, 66 kDa Bovine Serum Albumin, 45 kDa Ovalbumin, 36 kDa Glyceraldehyde 3-phosphate Dehydrogenase, 24 kDa trypsinogen, 20.1 kDa trypsin inhibitors and 14.2 kDa α-lactalbumin). After running SDS–PAGE gels, the protein bands were excised and in-gel trypsin digestion was performed folowed by ESI-QUAD-TOF Mass Spectrometry and Mascot (Matrix Science, UK) search engine against the NCBI NR database restricted to the taxa Snakes. The N-terminal amino acids sequence similarity was analyzed by alignment using BLAST (Altschul et al, 1997). The fibrinogenolytic activity (Edgar & Prentice, 1973) was assessed by SDS-PAGE after TLEs incubation with bovine fibrinogen releasing fibrinopeptides. The coagulant activity was performed qualitatively by evaluating the coagulation of human plasma in vitro, according Theakston & Reid, 1983.

TLEs from *B. moojeni* and *B. alternatus* were isolated in a faster than usual, two chromatographic steps procedure, with good protein yield. All TLEs shown high purity degree, molecular masses of 30-40 kDa and their partial sequences shared high identity with serine proteinases from other *Bothrops* snake venoms such as venom plasminogen activator, platelet-aggregating enzyme, SVSPs and batroxobin. The rapid purification and good yield of TLEs from *B. moojeni* and *B. alternatus* should provide new enzymes for improving the CEVAP’s heterologous fibrin sealant.

## RESULTS

1. Isolation of TLE serine proteinases from *Bothrops* snake venoms.
2. Yield of the purification process
3. Peptides identification by mass spectrometry
4. Serine Proteinases N-terminal amino acids sequencing
5. Multiple sequence alignments of TLEs N-terminal amino acid sequences
6. Fibrinogenolitic activity assay

## DISCUSSION

Snake venom serine proteinases (SVSPs) are present in several families, such as, Elapidae, Colubridae, Viperidae, and in particular, in the snake venom composition of a large number of *Bothrops* genus species. SVSPs demonstrate high specificity by the substrate, being capable of converting fibrinogen into fibrin (hence called thrombin-like enzymes) (Huang *et* al., 1999; Matsui *et* al., 1998), induce platelet aggregation (Basheer *et* al., 1995; Serrano *et* al., 1995), activate the X factor (Hofmann *et* al., 1983), increased capillary permeability (Sugihara *et* al., 1980), release bradykinin from the kininogen (Nikai *et* al., 1998; Serrano *et* al., 1998) and activate prothrombin (Kitano *et* al., 2013) among other activities. Between all activities produced by snake venom TLEs, the target of our bio-prospecting research is fibrinogenolitic activity (capacity of converting fibrinogen into fibrin) exerted by bothropic TLEs studied as a means to obtain information about future candidates for the composition of the fibrin heterologous sealant developed by CEVAP.

Several serine proteinases from *Bothrops* snakes venom have been purified and characterized, such as Bhalternin and Balterobin, wich have been isolated from *Bothrops alternatus* venom (Costa *et* al., 2010; Smolka *et* al., 1998), MSP 1, MSP 2, MMO3 and Batroxobin, isolated from *Bothrops moojeni* venom (Oliveira *et* al., 1999; Serrano *et* al., 1993; Stocker and Barlow, 1976), serine proteinases also have been identified in the venoms of *Bothrops jararacussu* (Bortoleto *et* al., 2002; Hill-Eubanks *et* al., 1989), *Bothrops atrox* (Itoh *et* al., 1987; Kirby *et* al., 1979; Petretski *et* al., 2000), *Bothrops jararaca* (Mandelbaum and Henriques, 1964; Nishida *et* al., 1994; Serrano *et* al., 1995), BpSP-I from *B. pauloensis* (Costa *et* al.,2009) and through a combination of three chromatographic steps, serine proteinases from *B. moojeni* and *B. alternatus* venoms have been isolated by Oliveira *et* al. in 2013 We presented in this work, a protocol to obtain serine proteinases from *B. alternatus* and *B. moojeni* venoms with high degree of purity, as showed by LC-MS and SDS-PAGE, suitable for structural and other biophysical and biochemical studies and clinical use.

All isolated serine proteinases, showed relative molecular masses, observed in the SDS-PAGE electrophoretic profile, between 30-40 kDa. Many serine proteinases of molecular masses of about 30 kDa share significant homology. Peptides data identified by mass spectrometry showed their partial sequences sharing high identity with other *Bothrops* SVSPs. The N-terminal analysis of purified serinoproteases to *B. alternatus* and *B. moojeni* and multiple alignment with other snake venoms TLEs (SVSP=Snake Venom Serine Proteinase; batroxobin=TLE - *B. atrox*; bhalternin=TLE - *B. alternatus*; asperase=TLE - *B. asper*; leucurobin= TLE - *B. leucurus*, chitribrisin=TLE - *Trimeresurus albolabris*; bilineobin= TLE - *Agkistrodon bilineatus*; stejnefibrase-2= Beta-fibrinogenase (TLE) - *Trimeresurus stejnegeri*; alpha-fibrinogenase= TLE - *Macrovipera lebetina*; BpirSP27= TLE – *B. pirajai*; saxthrombin= TLE - *Gloydius saxatilis*; cerastocytin= TLE - *Cerastes cerastes*; afaacytin= SVSP - *Cerastes cerastes*; cerastotin= TLE - *Cerastes cerastes*; venombin= TLE – *Agkistrodon controtrix contortrix*), demonstrated a high identity of the N-terminal sequence between them.

The SVSPs share among themselves a degree of similarity in amino acid sequence of about 65% on average (Oliveira *et* al., 2013). The high identity (67 to 92%) of purified serine proteinases samples from *B. alternatus* snake venom with fibrinogen clotting enzyme Bhalternin (Costa *et* al., 2010) obtained by the alignment of the N-terminal sequences(Figure 2), together with the elution profile analysis in RP-HPLC and electrophoretic profile (Figure1), leads us to the conclusion that these samples are the same TLE bhalternin, purified until then in three chromatographic steps.

**Figure 1.**
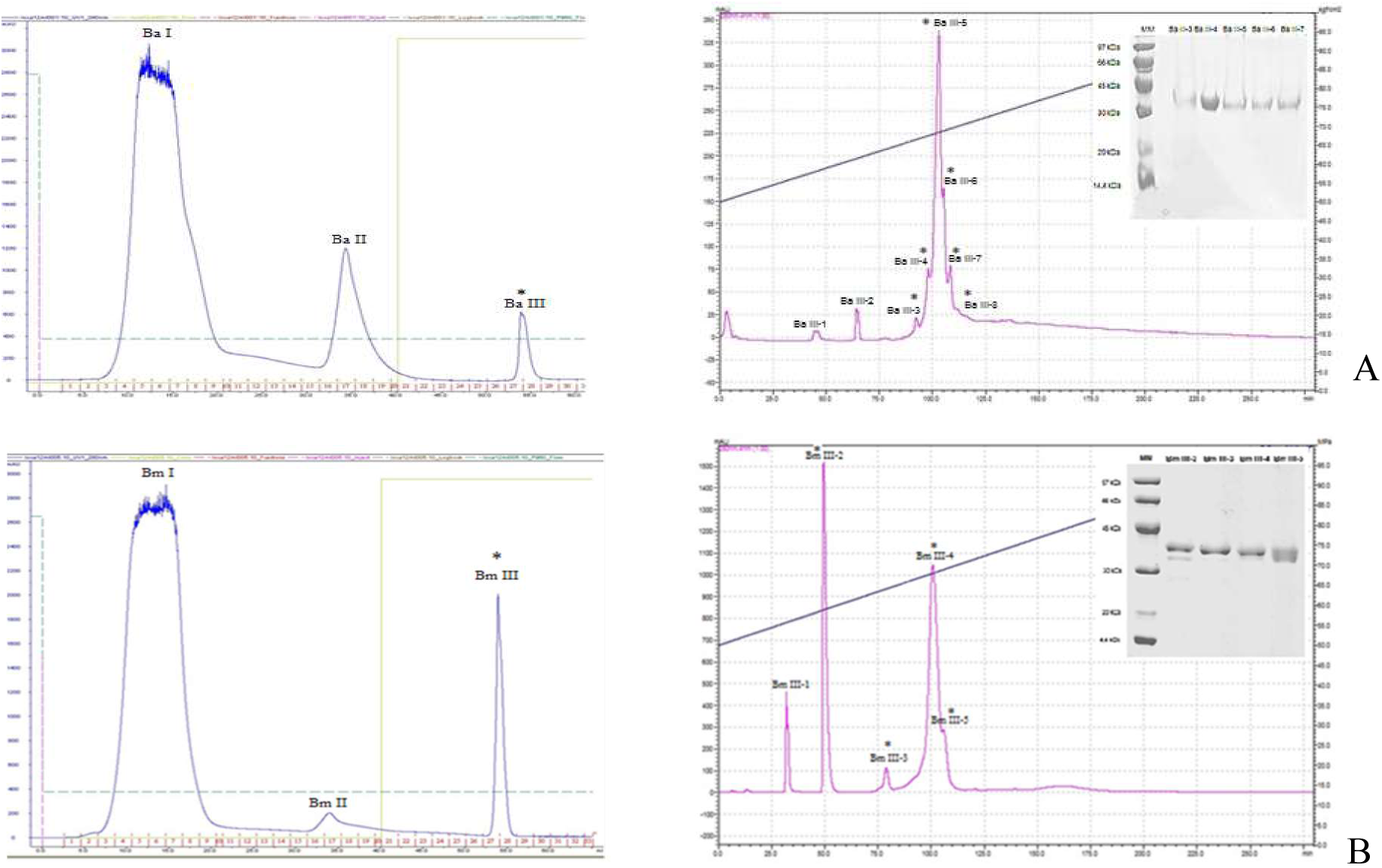
500 mg of crude venom of each *Bothrops* snake venom were applied to a Benzamidine-Seharose 6B affinity column. The first column figures represents the chromatograms obtained in this experimental phase, where the serine proteinases were located in peak III. Subsequently, the fraction III was subjected to RP-HPLC (second column) in a Shimadzu C18 column, revealing new fractions. The fractions marked with **\*** showed coagulant activity. **Insert:** Electrophoretic profiles on SDS-PAGE (13%) under denaturing and reducing conditions. **A**-*B. alternatus*; **B**-*B. moojeni*.

**Figure 2.**
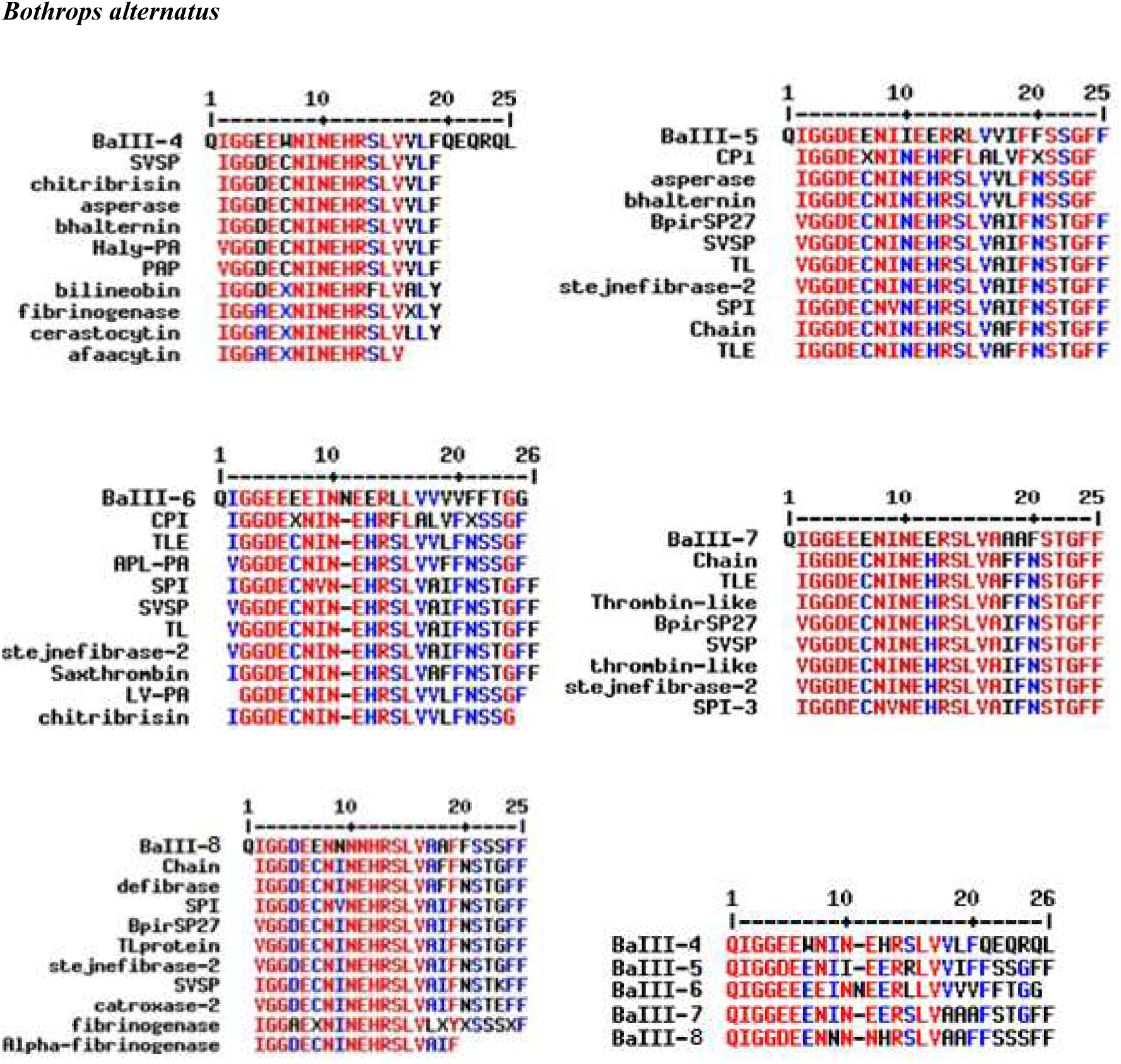
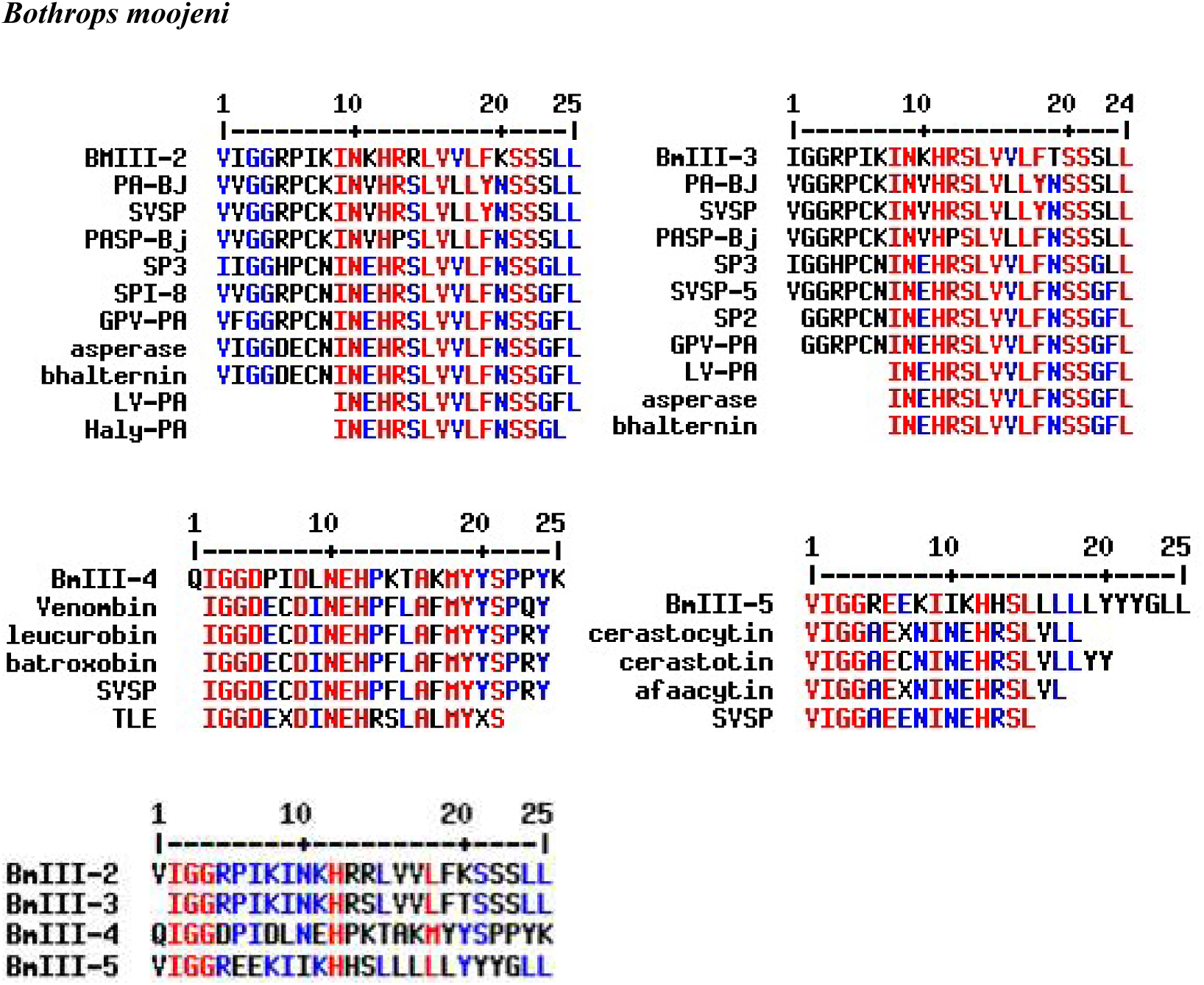
Multiple alignments of N-terminal amino acids sequences of thrombin-like serine proteinases (TLSP) samples from *B. alternatus* and *B. moojeni* venoms, obtained by the Edman degradation method. The N-terminal sequences of other SVSP were obtained from the BLAST-p database and aligned with the program MultAlin. SVSP=Snake Venom Serine Proteinase; batroxobin=TLE - *B. atrox*; bhalternin=TLE - *B. alternatus*; asperase=TLE - *B. asper*; leucurobin= TLE - *B. leucurus*; PA-BJ= Platelet-aggregating proteinase - *B. jararaca*; chitribrisin=TLE - *Trimeresurus albolabris*; Haly-PA= Venom plasminogen activator – *Gloydius brevicaudus*; bilineobin= TLE - *Agkistrodon bilineatus*; stejnefibrase-2= Beta-fibrinogenase (TLE) - *Trimeresurus stejnegeri*; catroxase-2= SVSP – *Crotalus atrox*; alpha-fibrinogenase= TLE - *Macrovipera lebetina*; BpirSP27= TLE – *B. pirajai*; saxthrombin= TLE - *Gloydius saxatilis*; cerastocytin= TLE - *Cerastes cerastes*; afaacytin= SVSP - *Cerastes cerastes*; cerastotin= TLE - *Cerastes cerastes*; venombin= TLE – *Agkistrodon controtrix contortrix*; GPV-PA= Venom plasminogen activator - *Trimeresurus albolabris*.

**Figure 3.**
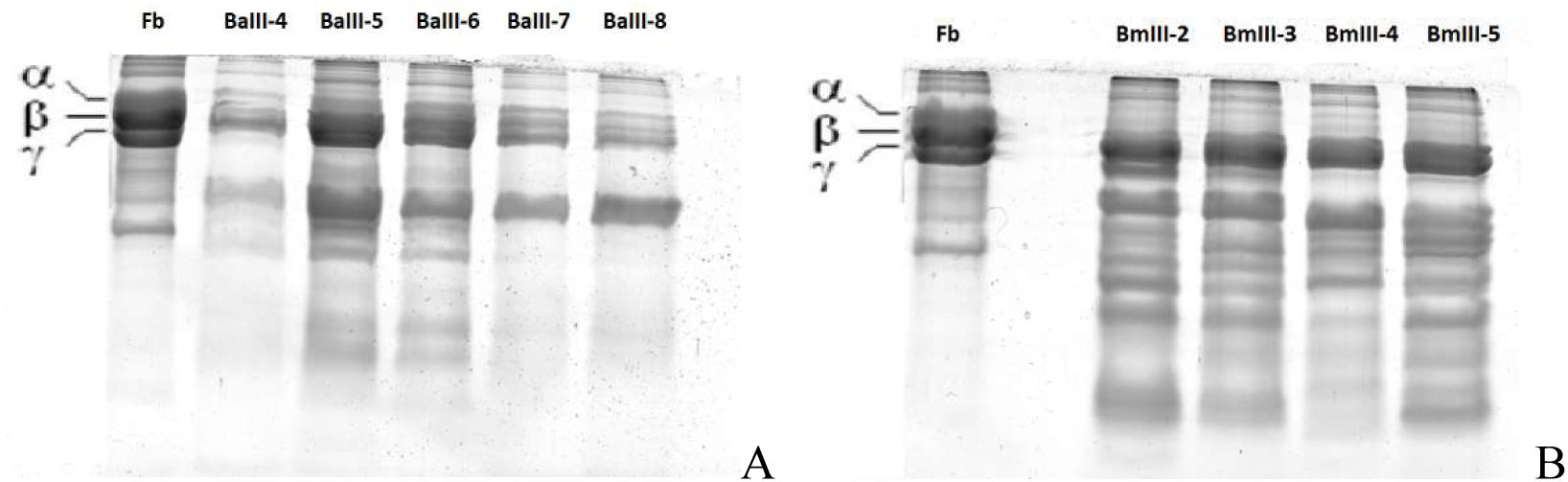
Analysis of bovine fibrinogen (Fb) degradation by TLE serine proteinases (10 µg) obtained from *B. alternatus* (A) and *B. moojeni* (B) venoms. The degradation of bovine fibrinogen (30 µg) was analyzed in 13% SDS-PAGE.

According to their specificity for cleaving fibrinogen chains, the serine proteinases have been classified as α, β and γ-fibrinogenases. SVSPs preferentially cleave the Bb-chain (Herzig et al., 1970; Markland, 1998; Menaldo et al., 2012; Vieira et al., 2004), however, some of them such as Batroxobin from *B. atrox*, Bothrombin from *B. jararaca* and Bhalternin from *B. alternatus* (Stocker and Barlow, 1976; Costa *et* al., 2010; Sant’Ana *et* al., 2008) cleaves the Aa-chain, being considered α-fibrinogenases, classical enzymes studied in basic and clinical research. The fibrinogenolitic (Fb) activity test performed with samples of thrombin-like serine proteinases obtained from *B. alternatus* and *B. moojeni* venoms demonstrate what this enzymes can be considered α-fibrinogenases, mainly due to the fact hydrolyze the Aα chain, and the TLE from *B. moojeni* showing greater activity than that from *B. alternatus*.

The knowledge about the components constituents of snake venoms has a great importance in the understanding of the venom as a whole. To understand the extreme complexity of these venoms, it is necessary to isolate these constituents and identify their individual properties. From this identification, we can search how to combat or explore the pharmacological effects caused by specific components of the venom.

## CONCLUSION

The purification process developed in this study is faster and more economical compared to the other existing methods, since it uses only two chromatographic steps for obtaining the serine proteinases of *B. alternatus* and *B. moojeni*. All isolated TLEs showed molecular mass of 30-40 kDa, and their partial sequences sharing high identity with other SVSPs, such as venom plasminogen activator, platelet-aggregating enzyme and lectins, according Mascot database search. Individual ions scores > 34 in table 2 indicate identity or extensive homology (p<0.05). The faster purification and good yield of TLEs from *B. moojeni* and *B. alternatus* should provide new knowledge about these enzymes. This study aimed to find alternatives to the thrombin-like enzyme from *Crotalus durissus terrificus* to produce the fibrin sealant and this job indicated several candidates to this replacement.

**Table 1.** *Bothrops* snakes crude venom total mass applied to the affinity chromatographic column, obtained mass of each venom fraction containing serine protease and the yield in the isolation process.

| Venom | Total venom mass applied (mg) to affinity chromatography | Protein dosage (mg) of serine proteinase-containing fractions after affinity chromatography | Yield (%) of serine proteinase-containing fractions isolated after affinity chromatography | Protein mass (mg) of serine proteinase-containing fractions isolated after RP-HPLC | Yield (%) of serine proteinase-containing fractions isolated after RP-HPLC | Fractions purification (n times) |
| --- | --- | --- | --- | --- | --- | --- |
| *Bothrops alternatus* | 500 | 14,6 | 2,92 | BaIII-4= 1,04 | 7,1 | 480 |
|  |  |  |  | BaIII-5= 5,32 | 36,4 | 94 |
|  |  |  |  | BaIII-6= 0,82 | 5,6 | 609 |
|  |  |  |  | BaIII-7= 1,01 | 6,8 | 495 |
|  |  |  |  | BaIII-8= 0,32 | 2,2 | 1562 |
| *Bothrops moojeni* | 500 | 32,0 | 6,4 | BmIII-2= 10,0 | 31,2 | 50 |
|  |  |  |  | BmIII-3= 1,98 | 6,2 | 252 |
|  |  |  |  | BmIII-4= 16,5 | 51,5 | 30 |
|  |  |  |  | BmIII-5= 2,76 | 8,6 | 181 |

**Table 2.**
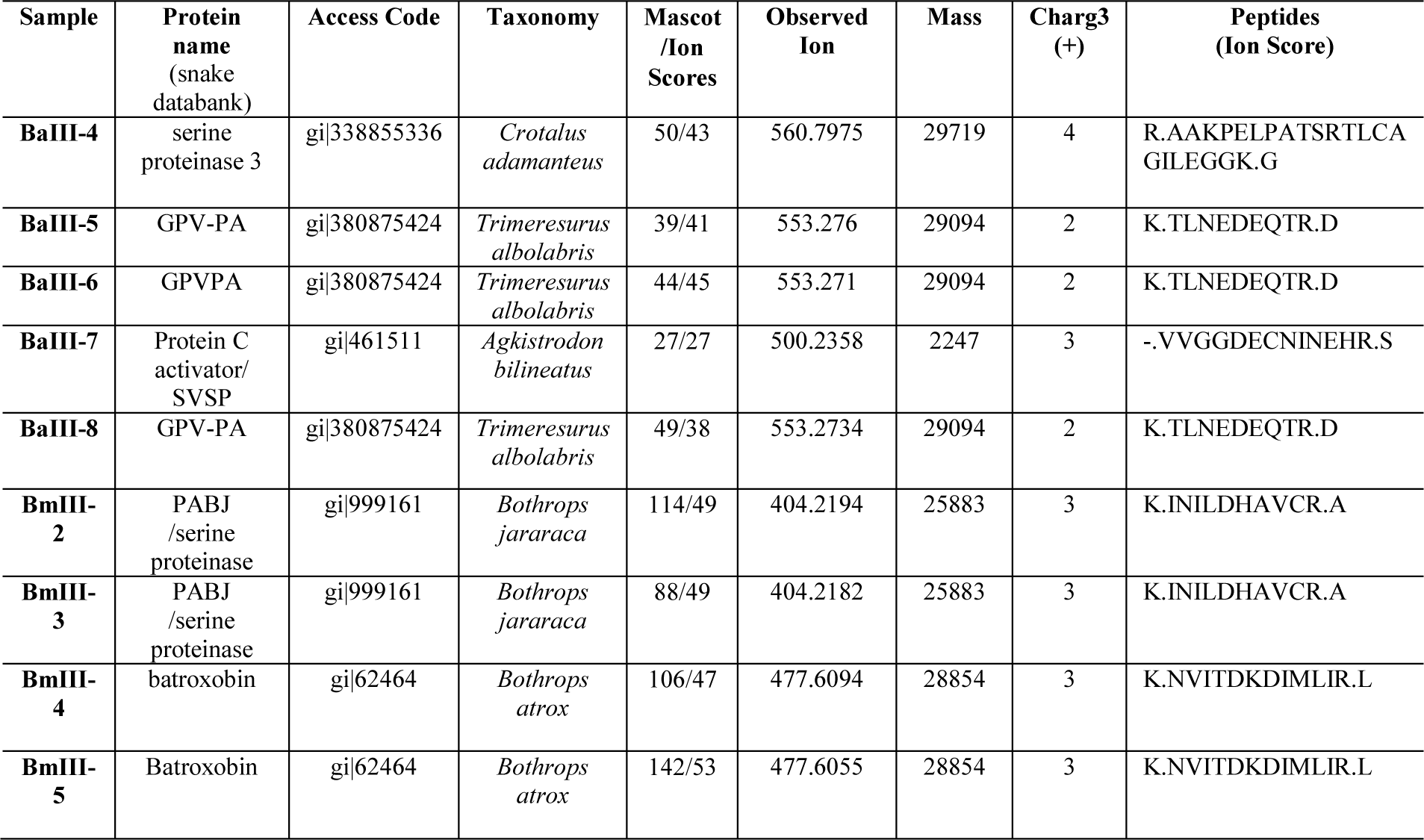
Data of peptides identified by mass spectrometry

**Table 3.**
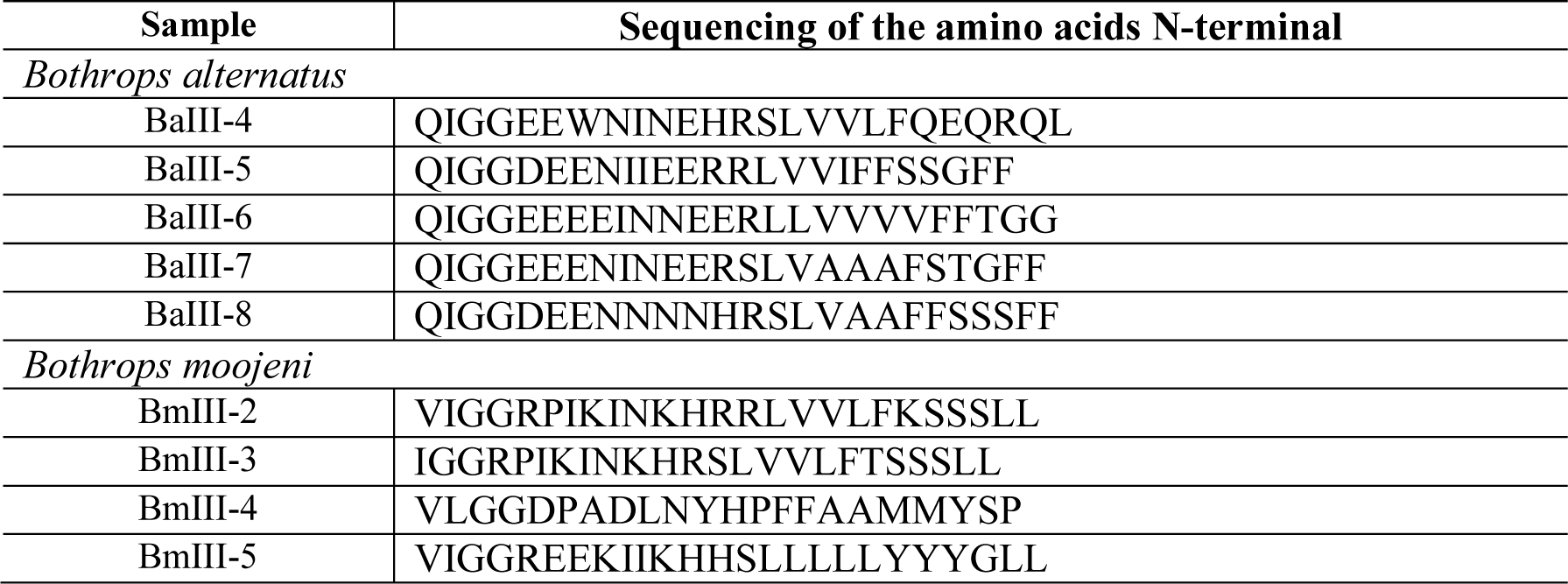
Sequencing of the amino acids N-terminal portion of TLEs purified from *B. alternatus* e *B. moojeni* venoms

